# Novel insights into the molecular heterogeneity of hepatocellular carcinoma

**DOI:** 10.1101/101766

**Authors:** Juan Jovel, Zhen Lin, Sandra O’keefe, Steven Willows, Weiwei Wang, Guangzhi Zhang, Jordan Patterson, David J. Kelvin, Gane Ka-Shu Wong, Andrew L. Mason

**Author notes:** These authors contributed equally to this work (Juan Jovel, Zhen Lin). Corresponding author: G.K. Wong, 4-126 Katz Group Centre for Pharmacy and Health Research, University of Alberta, Edmonton AB, T6G 2E1. A.L. Mason, 7-142 Katz Group Centre for Pharmacy and Health Research, University of Alberta, Edmonton AB, T6G 2E1. **Competing interest**: The authors declare that no competing interest exists.

## Abstract

Hepatocellular carcinoma (HCC) is influenced by numerous factors, which results in diverse genetic, epigenetic and transcriptional scenarios, thus posing obvious challenges for disease management. We scrutinized the molecular heterogeneity of HCC with a multi-omics approach in two small cohorts of resected and explanted livers. Whole-genome transcriptomics was conducted, including polyadenylated transcripts and micro (mi)-RNAs. Copy number variants (CNV) were inferred from whole genome low-pass sequencing data. Fifty-six cancer-related genes were screened using an oncology panel assay. HCC was associated with a dramatic transcriptional deregulation of hundreds of protein-coding genes suggesting downregulation of drugs catabolism, induction of inflammatory responses, and increased cell proliferation in resected livers. Moreover, several long non-coding RNAs and miRNAs not reported previously in the context of HCC were found deregulated. In explanted livers, downregulation of genes involved in energy-producing processes and upregulation of genes aiding in glycolysis were detected. Numerous CNV events were observed, with conspicuous hotspots on chromosomes 1 and 17. Amplifications were more common than deletions, and spanned regions containing genes potentially involved in tumorigenesis. CSF1R, FGFR3, FLT3, NPM1, PDGFRA, PTEN, SMO and TP53 were mutated in all tumors, while other 26 cancer-related genes were mutated with variable penetrance. Our results highlight a remarkable molecular heterogeneity between HCC tumors and reinforce the notion that precision medicine approaches are urgently needed for cancer treatment. We expect that our results will serve as a valuable dataset that will generate hypotheses for us or other researchers to evaluate to ultimately improve our understanding of HCC biology.

## Introduction

Hepatocellular carcinoma (HCC) is a lethal neoplasm, often secondary to chronic liver diseases induced by infections of hepatitis virus B (HBV) [1] and C (HCV) [2], exposure to aflatoxin B1 from Aspergillus [3], alcohol abuse [4], or from non-alcoholic steatohepatitis [5,6]. The disease is distributed worldwide [7] but its incidence is particularly high in Southeast Asia and the Sub-Saharan Africa [8–11], likely due to the high infection rate of HBV and exposure to aflatoxin B1 from contaminated grains; nonetheless, HCC occurrence is also growing in Western countries [7]. Despite considerable advances on HCC diagnosis and treatment, the proportion of resectable HCC tumors remains low, essentially due to lack of effective early diagnosis approaches, and recurrence after resection is high [12].

Broadly speaking, the natural history of HCC can be divided into i) molecular, ii) preclinical and iii) clinical or symptomatic phases [13]. The molecular phase refers to tumor biogenesis, which regularly follows pre-neoplastic stages characterized by chronic hepatitis and cirrhosis, and entails transformation of hepatocytes, biliary epithelial or stem cells [14,15]. Genetic alterations in differentiated cells often evolve into highly proliferative and recalcitrant-to-apoptosis cells, while transformation of stem cells leads to aberrant cell differentiation [13]. Initial epigenetic alterations in transcription are due to exposure to high levels of growth factors and pro-inflammatory cytokines, and are followed by chromosomal rearrangements that result in amplification and losses of proto-oncogenes and tumor suppressors, respectively. HBV integrations also lacerate the genome creating genetic aberrations [16]. Moreover, both HBV and HCV have the potential to directly modulate pathways that contribute to hepatocyte transformation [17,18], to prompt re-expression of TERT –to surmount telomere shortening– and to induce chromosomal instability by disrupting mitosis checkpoints [19,20]. In the preclinical phase, the tumor may or may not be detectable, and symptoms are still not noticeable in the patient. Finally, the clinical phase is associated with symptoms and usually occurs when the tumor is between 4-8 cm [21].

Studying the natural history of cancer, i.e. its ontogenesis, is of paramount importance because it offers the possibility of intervening patients at the most potentially effective stage during disease development. A high resolution picture of HCC’s ontogenesis is being portrayed by the accumulation of high-throughput data on cancer genomics, epigenomics, transcriptomics, proteomics and metabolomics, and their integration into computational frameworks [22]. Although considerable progress has been achieved, our understanding of the molecular diversity of HCC is still in its infancy. Ultimately, it should result in improved diagnostic and management strategies, and so will provide the basis for effective surveillance programs.

Many diseases, but very especially cancer, are associated with aberrant genomic and transcriptional landscapes [23]. This extends beyond protein-coding genes into several classes of structurally and functionally different non-coding RNAs [24–26]. We undertook a multi-omics approach to investigate the molecular heterogeneity of HCC tumors at early (resected livers) and late (explanted livers) stages of development. Our results provide a series of novel insights into the molecular heterogeneity of hepatocellular carcinoma, which may be further investigated in search of biomarkers or therapeutic drug targets.

## Results

### RNAseq profiling of HCC in resected livers

We sequenced and analyzed the transcriptome of liver sections affected by hepatocellular carcinoma or sections in the same livers not affected by cancer, initially from patients at early stages of the disease, who were subjected to liver resection. Tumoral tissue or corresponding control tissues were collected from twelve patients, ranging from 45 to 77 years old, exhibiting diverse pathologies (see Supp. Table 1 for details). In addition to this cohort of resected samples, we reanalyzed RNAseq data of 50 paired tumor-control samples, from resected livers, from The Cancer Genome Atlas database [27] to explore the reproducibility of our own findings (Supp. Fig. 1). Henceforth we refer to such samples as the TCGA cohort.

**Table 1.** Cancer-related genes mutated in each of the analyzed samples. Numbers indicate the number of mutations detected per gene.

| Gene | 1T | 2T | 3T | 4T | 5T | 6T | 7T | 8T | 9T | 10T | 11T | 12T | 311T | 315T | 338T |
| --- | --- | --- | --- | --- | --- | --- | --- | --- | --- | --- | --- | --- | --- | --- | --- |
| ABL1 |  |  |  | 1 |  |  |  |  |  |  |  |  |  |  |  |
| AKT1 |  |  |  |  |  |  |  |  |  |  |  |  |  |  |  |
| ALK | 1 | 1 |  | 1 | 1 | 1 | 1 | 1 | 1 |  | 1 | 1 | 1 | 1 | 1 |
| APC | 1 | 1 | 1 | 1 | 1 | 1 | 1 | 1 | 1 |  |  | 1 |  |  | 1 |
| ATM |  |  |  | 1 |  |  |  |  | 1 |  |  |  |  |  |  |
| BRAF |  |  |  |  |  |  |  |  |  |  |  |  |  |  |  |
| CDH1 |  |  |  |  |  |  |  |  |  |  |  |  |  |  |  |
| CDKN2A |  |  |  |  |  |  |  |  |  |  |  |  |  |  |  |
| CSF1R | 2 | 2 | 2 | 2 | 2 | 2 | 2 | 2 | 2 | 2 | 2 | 2 | 2 | 2 | 2 |
| CTNNB1 | 1 |  |  |  | 1 |  | 1 | 2 | 1 |  |  | 2 |  |  |  |
| DDR2 |  |  |  |  |  |  |  |  |  |  |  |  |  |  |  |
| DNMT3A | 2 |  | 2 | 2 |  |  |  | 2 |  |  | 2 |  |  |  |  |
| EGFR | 1 | 1 |  | 3 | 2 | 2 | 3 | 1 | 2 |  | 3 | 1 | 2 | 4 | 1 |
| ERBB2 |  |  |  |  |  |  |  |  |  |  |  |  |  |  |  |
| ERBB4 | 1 |  | 1 |  |  | 1 |  |  | 1 | 1 | 1 | 1 |  |  | 1 |
| EZH2 |  |  |  |  |  |  |  |  |  |  |  |  |  |  |  |
| FBXW7 |  |  |  |  |  |  |  |  |  |  |  |  |  | 1 |  |
| FGFR1 |  | 1 |  |  |  |  |  |  |  |  | 3 | 2 |  |  |  |
| FGFR2 |  |  |  |  |  |  |  |  |  |  |  |  |  |  |  |
| FGFR3 | 3 | 2 | 1 | 3 | 3 | 2 | 3 | 4 | 2 | 2 | 2 | 2 | 1 | 2 | 4 |
| FLT3 | 2 | 1 | 1 | 1 | 1 | 1 | 1 | 1 | 1 | 2 | 1 | 1 | 1 | 1 | 1 |
| FOXL2 |  |  |  |  |  |  |  |  |  |  |  |  |  |  |  |
| GNA11 | 1 |  |  | 1 |  |  |  |  | 1 |  |  | 1 |  |  |  |
| GNAQ |  |  |  |  |  |  |  |  |  |  |  |  |  |  |  |
| GNAS |  |  |  |  |  |  |  |  |  |  | 1 |  |  |  |  |
| HNF1A | 1 |  |  |  |  |  | 1 |  |  | 1 | 1 |  |  | 1 | 1 |
| HRAS |  | 1 |  | 1 | 1 |  | 1 |  |  | 1 | 1 | 1 | 1 |  |  |
| IDH1 |  |  |  |  |  |  |  |  |  |  |  |  |  | 1 |  |
| IDH2 |  |  |  | 1 | 2 |  |  |  |  |  |  |  |  |  |  |
| JAK2 |  |  |  |  |  |  |  |  |  |  |  |  |  |  |  |
| JAK3 |  | 1 |  | 1 |  |  |  |  | 1 |  |  |  |  |  |  |
| KDR | 2 | 2 | 4 | 4 |  | 4 | 4 | 2 | 1 | 1 | 1 |  | 4 | 1 | 1 |
| KIT |  |  | 2 |  |  | 2 |  |  | 2 | 2 | 1 |  | 2 | 1 |  |
| KRAS |  |  |  |  |  |  |  | 1 |  |  |  |  |  |  |  |
| MAP2K1 |  |  |  |  |  |  |  |  |  |  |  |  |  |  |  |
| MET |  |  | 2 |  |  |  |  |  |  |  |  |  |  |  |  |
| MLH1 |  |  |  |  |  |  |  |  |  |  |  |  |  |  |  |
| MPL |  |  |  |  |  |  |  |  |  |  |  |  |  |  | 1 |
| MSH6 | 1 | 1 | 1 |  | 1 | 1 |  | 1 | 1 |  |  | 1 | 1 | 1 | 1 |
| NOTCH1 |  |  |  |  |  |  |  |  |  |  |  |  |  |  |  |
| NPM1 | 1 | 1 | 1 | 1 | 1 | 1 | 1 | 1 | 1 | 1 | 1 | 1 | 1 | 1 | 1 |
| NRAS |  |  |  |  |  |  |  |  |  |  |  |  |  |  |  |
| PDGFRA | 6 | 4 | 4 | 3 | 5 | 1 | 1 | 4 | 4 | 5 | 2 | 3 | 4 | 4 | 5 |
| PIK3CA |  |  |  |  |  |  |  | 2 |  |  |  |  |  |  | 1 |
| PTEN | 1 | 1 | 1 | 1 | 2 | 1 | 1 | 1 | 1 | 1 | 1 | 2 | 1 | 1 | 1 |
| PTPN11 |  |  |  |  |  |  |  | 1 |  |  |  |  |  |  |  |
| RB1 |  |  |  |  |  |  |  |  |  |  |  |  |  |  |  |
| RET | 2 | 4 |  | 2 | 2 |  | 3 | 1 | 3 |  | 1 |  | 1 | 3 | 1 |
| SKT11 |  |  |  |  |  |  |  |  |  |  |  |  |  |  |  |
| SMAD4 |  |  | 1 |  |  | 1 |  |  |  |  | 1 |  |  |  |  |
| SMARCB1 |  |  |  | 1 |  | 1 | 1 |  | 1 | 1 | 1 |  |  |  | 1 |
| SMO | 1 | 1 | 1 | 1 | 2 | 1 | 2 | 5 | 3 | 1 | 1 | 1 | 2 | 1 | 1 |
| SRC |  |  |  |  |  |  |  |  |  |  |  |  |  |  |  |
| TP53 | 4 | 4 | 3 | 4 | 5 | 5 | 3 | 3 | 3 | 4 | 3 | 4 | 5 | 4 | 4 |
| TSC1 |  |  |  |  |  |  |  |  | 1 |  |  |  |  |  |  |
| VHL |  |  |  |  |  |  |  |  |  |  |  |  |  |  |  |

**Figure 1.**
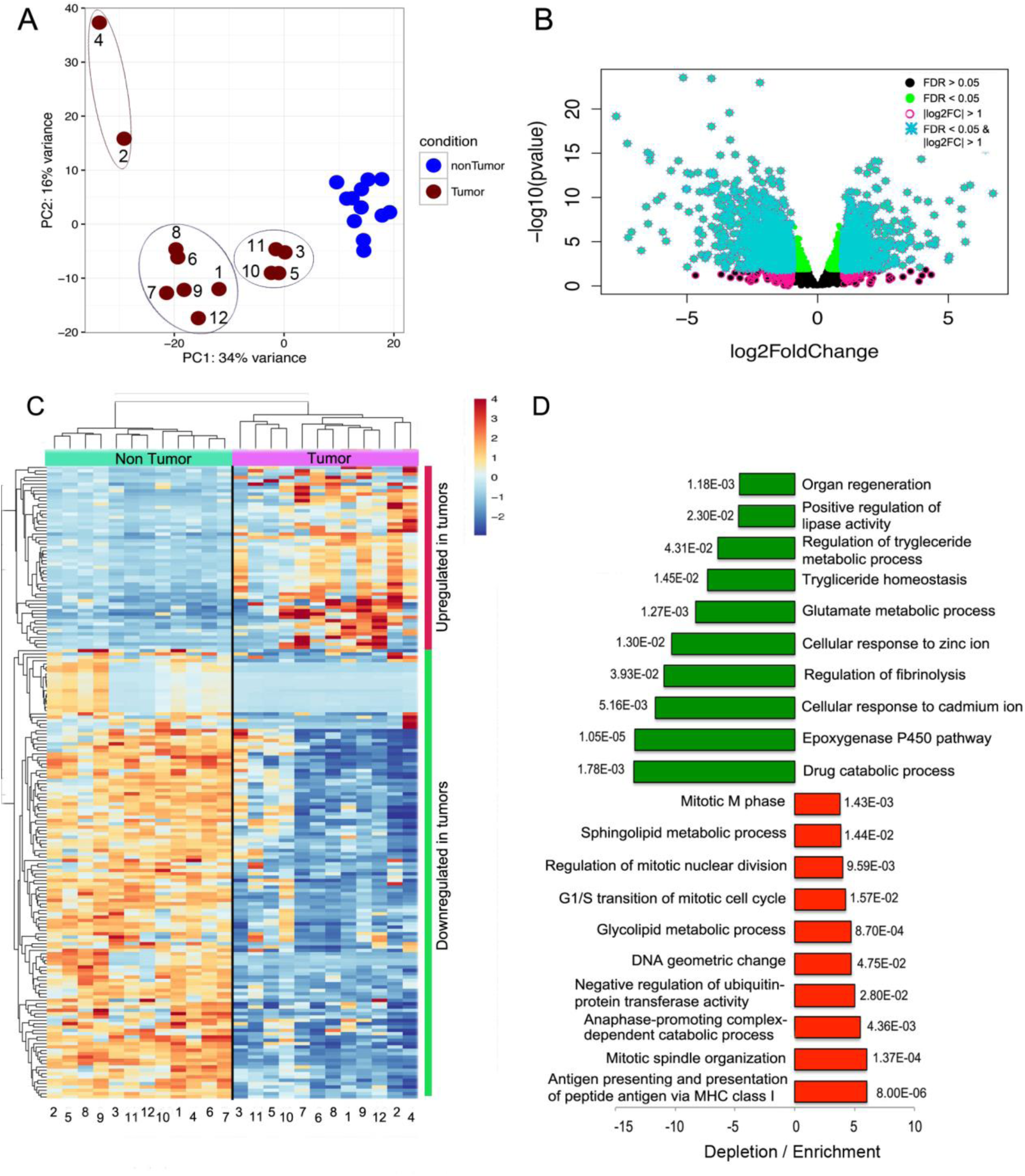
Dramatic transcriptional deregulation in HCC resected tumors hints at biogenesis of drug resistance and activation of inflammation. (**A**) Principal component analysis clearly separated tumor and non-tumor samples. While non-tumor samples clustered rightly, tumor samples dispersed along the two axis (components) forming apparent sub-clusters. (**B**) Classification of differentially expressed genes according to false discovery rate (FDR) and log2 fold change. Deregulated genes with a FDR < 0.05 (green dots) and log2 fold change > 1 were considered as truly differentially expressed in this study (turquoise asterisks). (**C**) Hierarchical clustering using the Z scores of the 200 deregulated genes that exhibited the largest variance also separated tumor and non-tumor samples into separate branches of a dendrogram, with larger distances in the tumor branches. Tumors were divided into the same sub-clusters illustrated in the PCA plot. (**D**) Gene ontology analyses were conducted in the Gene Ontology Consortium web portal (http://www.geneontology.org/). The ten most enriched or most depleted terms are presented along with the corresponding P values. In general, the magnitude of depletion is larger than the magnitude of enrichment. The most depleted terms were *Drug catabolicprocesses* and the *Epoxygenase P450 pathway* which are indicative of drug resistance emergence and activation of the immune system, respectively. Top enriched terms are related to DNA synthesis and mitosis, suggestive of increase cell proliferation in tumors.

In principal component analyses, non-tumor samples clustered relatively close to each other, while tumor samples distributed more dispersedly with two tumors showing considerable deviation from the rest of tumors (Fig. 1A; tumors 2 and 4). Tumors 3, 5, 10 and 11 formed a sub-cluster, while the rest of samples formed a less-compact subcluster (Fig. 1A). When a false discovery rate < 0.05 and a fold change > 2 were assumed as thresholds for statistical significance, 726 and 1,066 genes were found upregulated and downregulated, respectively in HCC samples (Supp. Table 2; Fig. 1B). To provide a more detailed perspective on gene deregulation, the Z scores of 200 deregulated genes exhibiting the largest fold change are depicted in Fig. 1C. In the upper part of the heatmap agglomerate many genes that are considerably upregulated in some tumors, but heterogeneity in their expression across tumors is notable. Downwards, a larger set of genes appears severely downregulated in tumors. In concordance with the separation observed in the PCA plot, some tumors, like 3, 11, 5 and 10, formed a separate clade in the upper dendrogram and tumors 2 and 4 also branched apart from the rest of tumors.

The magnitude of the downregulation events was greater than the one of upregulation events (Fig. 1B). For instance, 97, 25 and 14 genes were downregulated 10-, 25-, 50-fold or more, while only 36, 10 and 4 genes were upregulated 10-, 25-, 50-fold or more (Supp. Table 2). The same trend was also observed in the TCGA cohort (Supp. Table 6). Among the most downregulated genes were some encoding members of the type-C lectin domain family 4 (CLEC4M* and CLEC4G*), the cholinergic receptor nicotinic alpha 4 subunit (CHRNA4*), the urocanate hydratase 1 (UROC1*) and a member of the cytochrome P450 family 2 (CYP2A7*). Among the most upregulated genes were an aldehyde dehydrogenase (ALDH3A1), a glutamate transporter (SLC7A11*), an aldo/keto reductase (AKR1B10*), the lipocain2 (LPC2), also known as oncogene 24p3 [28], and the anillin ANLN* (Supp. Table 2). Genes marked with asterisks were also found deregulated in the TCGA cohort in an analogous way (Supp. Table 6 or [29]). Another interesting set of deregulated genes are different effectors of oxidative phosphorylation (Fig. 2 and Supp. Table 2), including subunits of the ATP synthase complex (MT-ATP6, MT-ATP8), subunits of the cytochrome C oxidase (MT-CO1, MT-CO2, MT-CO3), subunits of the ubiquinol cytochrome C reductase (MT-CYB), subunits of the NADH dehydrogenase (MT-ND1, MT-ND2, MT-ND3, MT-ND4, MT-ND4L, MT-ND5, MT-ND6), the mitochondrial ribosomal RNAs 12S and 16S (MT-RNR1, MT-RNR2), a tRNA (MT-TP), and many metallothioneins (Fig. 2 and Supp. Table 2).

**Figure 2.**
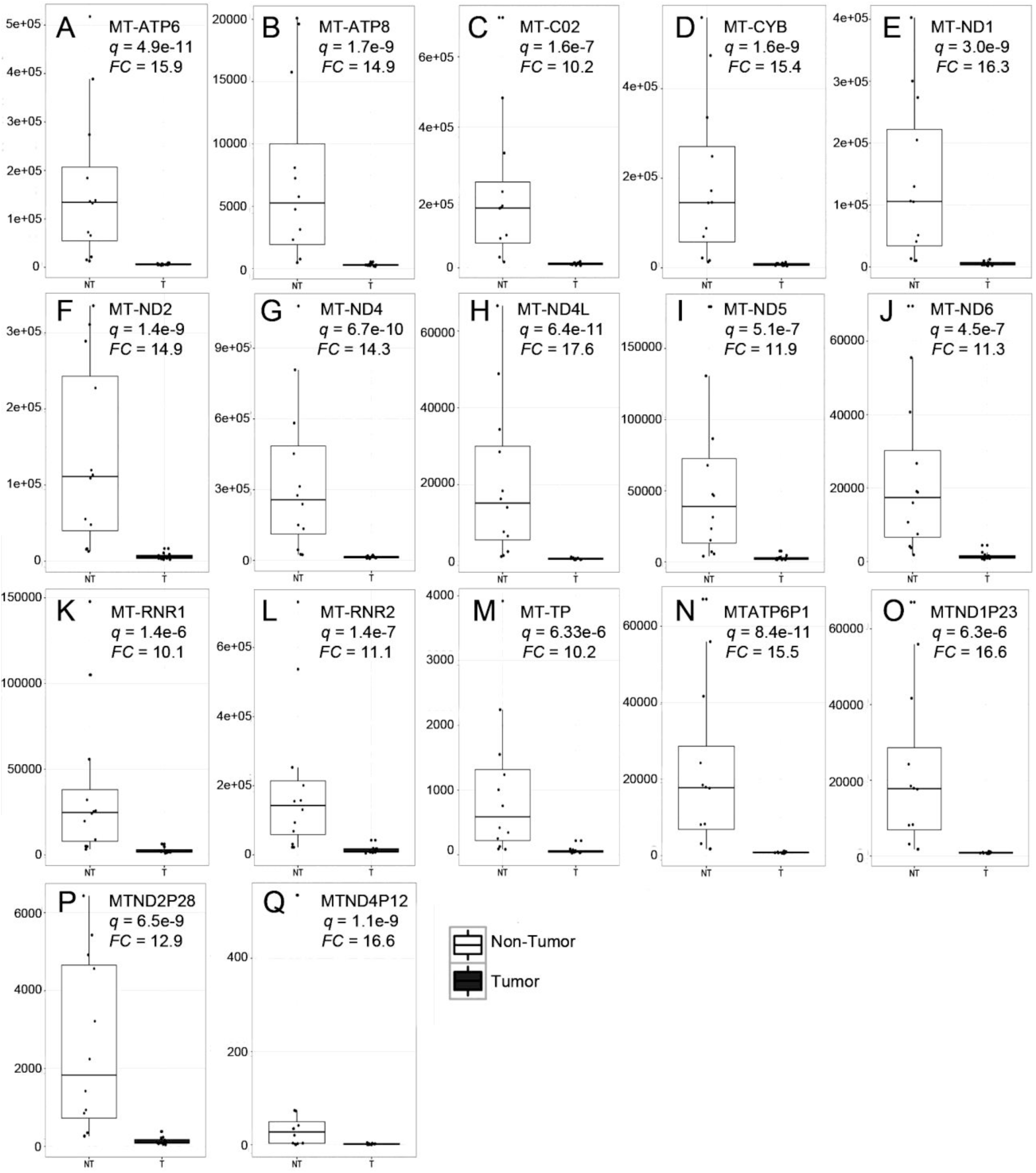
HCC is associated with a dramatic downregulation, but not shut-off, of enzymes involved in oxidative phosphorylation. Only genes that were found exclusively deregulated in our resected livers cohort, but neither in explanted livers, nor in the TCGA cohort, are presented here. Only genes that were found downregulated 10-fold or more are shown. Downregulated genes include subunits of the ATP synthase (**A, B**), a subunit of the cytochrome C oxidase (**C**), a subunit of the respiratory chain protein ubiquinol cytochrome C reductase (**D**), subunits of the NADH dehydrogenase (**E-J**), mitochondrial ribosomal RNAs (**K, L**), a mitochondrial tRNA (**M**) and pseudogenes for the mitochondrially-encoded NADH ubiquinone oxidoreductase core subunits (**N-Q**). The importance of pseudogenes on disease in currently under intense debate [76].

We also observed a substantial downregulation of many long non-coding RNAs (lncRNA) [26,30], with the exception of LINC00152 (aka CYTOR), which was found upregulated 3.6-fold (Fig. 3). Especially interesting are LINC01093 and LINC01595, which were found 30- and 10-fold downregulated. However, most lncRNAs were downregulated in some, but not all, tumors.

**Figure 3.**
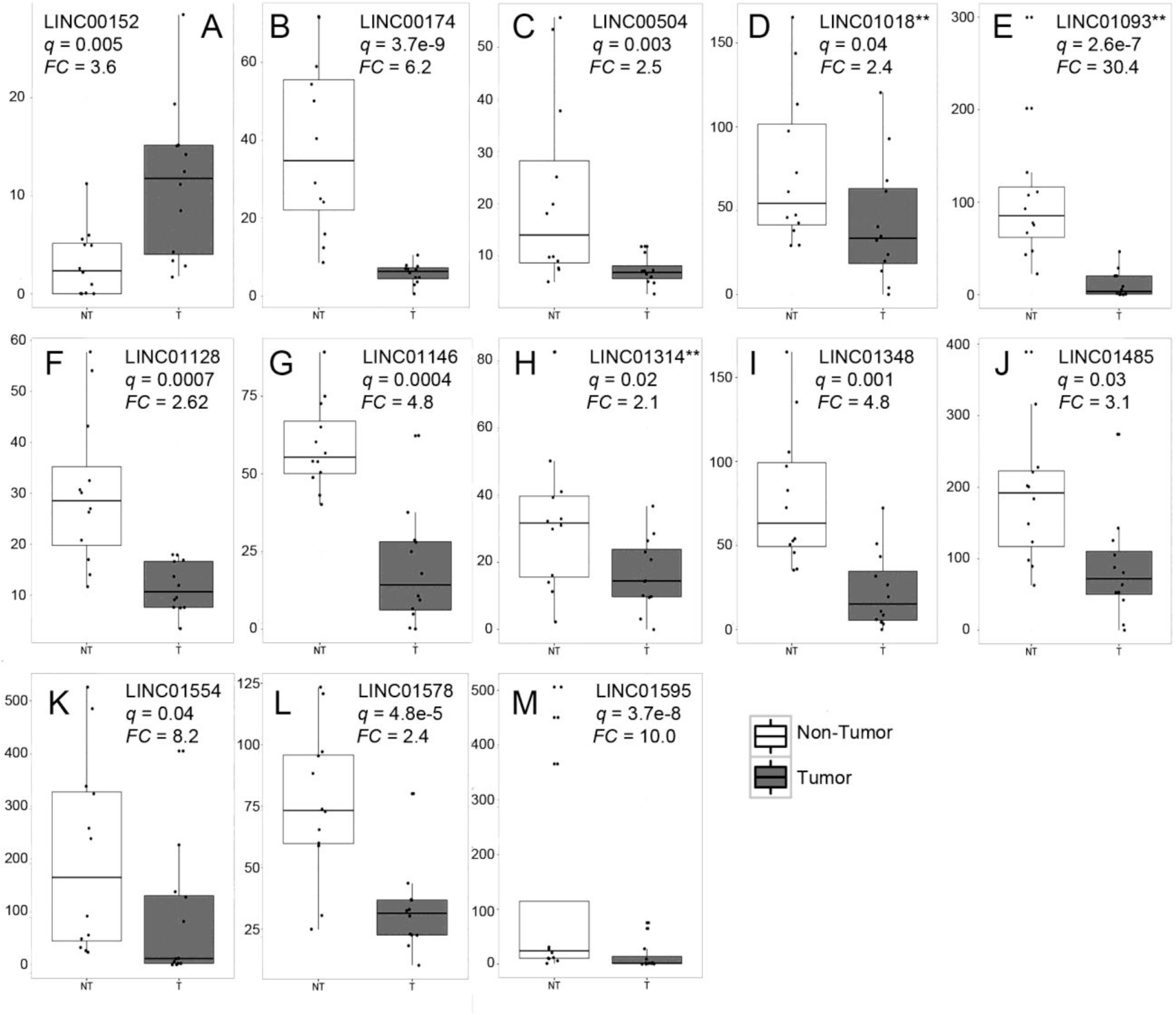
Many long non-coding RNAs (lncRNAs) are deregulated in resected HCC samples. (**A**)LINC00152 was the only lncRNA found upregulated in tumor tissue. All other lncRNAs found deregulated were downregulated in tumor samples (**B-M**). ** lncRNAs that were found deregulated in the resected and explanted HCC cohorts.

We then conducted gene ontology (GO) analyses [31]. The main biological process deregulated was *Drug catabolic processes* (13.5-fold decrease; p=1.8e-3; Fig. 1D), which was also found downregulated in the TCGA cohort (Supp. Fig. 2), this is reflected in the large number of severely downregulated genes belonging to the Cytochrome P450 superfamily of enzymes (Supp. Table 2; Supp. Table 6; Supp. Fig. 3) and to a lesser extent glutathione transferase genes (Supp. Table 2 and Supp. Table 6). Downregulation of drug catabolic process is associated with cancer drug resistance. Another GO term significantly downregulated was the *Epoxygenase P450 pathway* (Fig. 1D), which is also mediated by CYP enzymes (Supp. Fig. 3). In addition to xenobiotics, CYP enzymes subfamilies CYP2C and CYP2J also metabolize arachidonic acid to metabolically active epoxyeicosatrienoic acids (EETs), which are transient signaling molecules, whose metabolism inversely correlates with activation of the immune system and inflammation [32]. Accordingly, we found that CYP2AC8 and CYP2C9 were downregulated in cancer tissue 7.1- ad 5.2-fold, respectively (Supp. Fig. 3G and 3H). Among the predominantly upregulated pathways were those related to mitosis, lipid processing and antigen presentation (Fig. 1D).

### RNAseq profiling of HCC in explanted livers

To gain insights into the transcriptional deregulations associated with late stages of HCC, we analyzed cancer and control tissue from explanted livers from three patients affected by metastatic HCC (Supp. Table 1). The transcriptional profiles in non-tumor tissues were more similar among each other than were to their cancer counterparts (Fig. 4A), further highlighting transcriptional heterogeneity in cancer samples. A larger number of differentially expressed genes were downregulated (574 downregulated versus 343 upregulated) and the magnitude in fold-change was larger for downregulated genes (Fig. 4B and Supp. Table 3). RNA expression profile in tumors was heterogeneous across the 200 most variable deregulated genes (Fig. 4C). The repertoire of deregulated genes was substantially different from genes deregulated in resected livers. Indeed, only 69 genes were deregulated in both resected and explanted HCC tumors with the same polarity (Supp. Table 8 and Supp. Table 9). In gene ontology analysis, the most depleted term was *Short-chain fatty acid catabolism*; short-chain fatty acids, especially butyrate, are associated with cellular homeostasis [33]. Other GO terms associated with the production of energy through cellular respiration, including the tricarboxylic acid cycle and consequently oxidative phosphorylation and terms related to production of amino acids were also found depleted (Fig. 4E). Some terms were moderately upregulated including *Cellular response to decreased oxygen levels* and *Response to hypoxia*.

**Figure 4.**
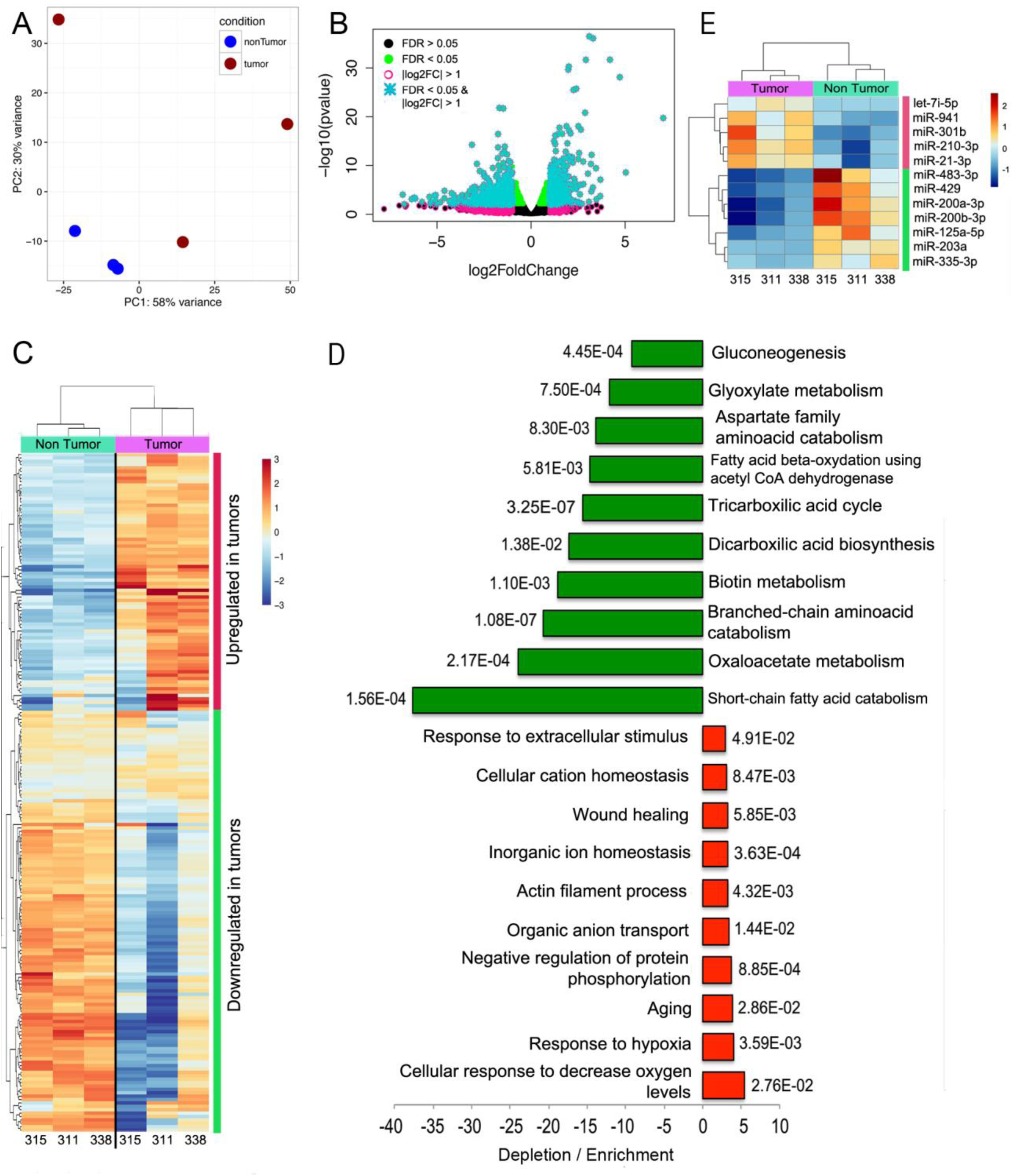
Transcriptional deregulation in HCC from explanted end-stage livers suggest general reduction of energetic processes. (**A**) Principal component analysis, as in the case of resected tumors, separated tumor from non-tumor samples, with the former ones exhibiting a more disperse distribution along the two principal components. One of the tumors (315) was located in close proximity to non-tumor samples. (**B**) Classification of differentially expressed genes according to false discovery rate (FDR) and log2 fold change. Deregulated genes with a FDR < 0.05 (green dots) and log2 fold change > 1 were considered as truly differentially expressed in this study (turquoise asterisks). (**C**) Hierarchical clustering using the Z scores of the 200 deregulated genes that exhibited the largest variance also separated tumor and non-tumor samples into separate branches of a dendrogram, with larger distances in the tumor branches. (**D**) Gene ontology analyses were conducted in the Gene Ontology Consortium web portal (http://www.geneontology.org/). The ten most enriched or most depleted terms are presented along with the corresponding P values. In general, the magnitude of depletion is larger than the magnitude of enrichment. (**E**) Few miRNAs were found deregulated in tumor samples, most of which have already been characterized in the context of HCC.

### microRNAseq profiling of HCC in resected and explanted livers

We explored the miRNAs profile in tumor and non-tumor tissues from all samples described above, including the TCGA cohort. Using the same thresholds (FDR < 0.05; fold change > 2) we found 30 and 44 miRNAs significantly upregulated or downregulated in our resected livers samples, respectively (Supp. Table 4; Fig. 5B). Principal component analysis, as in the case of the RNAseq data, also showed tightly clustered non-tumor samples and more dispersedly distributed tumor samples (Fig. 5A). Some tumor samples (i.e. 7 and 8) clearly separated from the rest. We noticed that the so-called 19C miRNA oncogenic cluster [34] was dramatically upregulated in tumors 7 and 8 (Supp. Fig. 4), and this may explain the location of such tumors on the first principal component axis (Fig. 5A). Some other samples, like 1, 3, 5, 10, 11 and 12 or 2, 4, 6 and 9 also formed sub-clusters (Fig. 4A). When the Z scores of the top 96 deregulated miRNAs were subjected to hierarchical clustering, rather discrete blocks of miRNAs that were overexpressed or underexpressed in tumors are clearly delineated (Fig. 4C). Samples 2 and 4 branched apart from the rest of tumor samples in the cladogram, and formed their own branch closer to the non-tumor samples.

**Figure 5.**
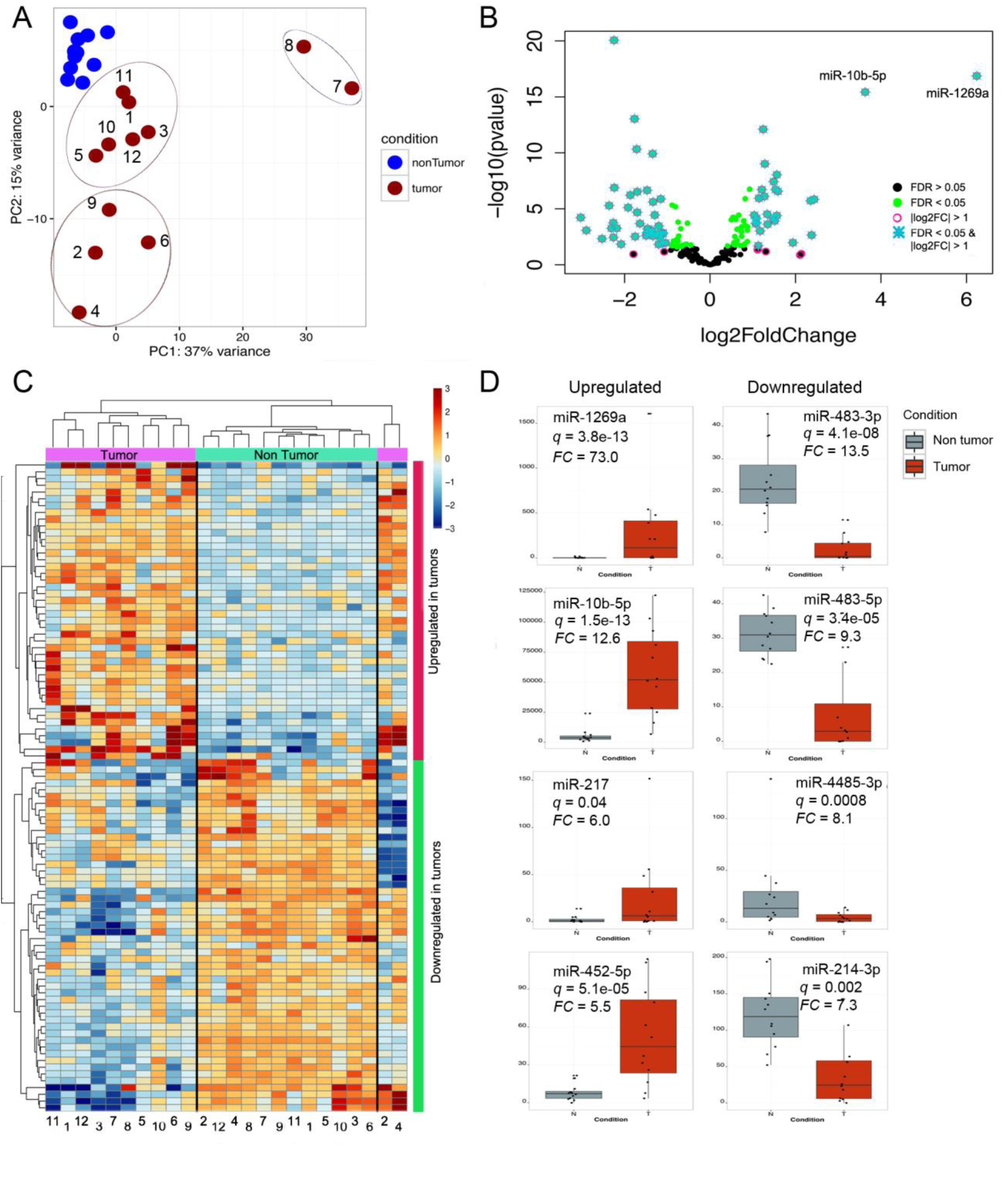
miRNA profiling reveals deregulated miRNAs that have not been previously characterized in the context of HCC. (**A**) Principal component analysis of the miRNA expression data clearly separated tumor from non-tumor samples. As in the case of RNAseq data, non-tumor samples formed a much more compact cluster than their tumor counterpart. Tumors 7 and 8 were considerably different from the rest of tumors; tumors 1,3,5,10,11, and 12 formed an apparent sub-cluster, as did tumors 2,4,6 and 9. (**B**) Classification of differentially expressed miRNAs according to false discovery rate (FDR) and log2 fold change. Deregulated genes with a FDR < 0.05 (green dots) and log2 fold change > 1 were considered as truly differentially expressed in this study (turquoise asterisks). miR-10b-5p and miR-1269a were substantially more deregulated that the rest of miRNAs and for this reason are explicitly shown here. (**C**) Hierarchical clustering of the Z scores of the 96 deregulated miRNAs exhibiting the largest variance separated most tumor from non-tumor samples, with the exception of tumors 2 and 4, which formed a branch apart from the rest of tumors, and closer to non-tumor samples. (**D**) The top four upregulated or downregulated miRNAs are shown.

The most upregulated miRNAs were miR-1269a (73-fold), miR-10b-5p (12.6-fold), miR-217 (6-fold) and miR-452-5p (5.5-fold). The most downregulated miRNAs were the two isoforms of miR-483 (3p=13.5-fold and 5p=9.3-fold), miR-4485-3p (8.1-fold), and miR-214-3p (7.3-fold). Except for miR-483-3p and miR-4485-3p all the above-mentioned miRNAs were also found deregulated, with the same polarity, in the TCGA cohort (Supp. Table 7). In explanted livers, perhaps because of the reduced number of samples, only five and seven upregulated and downregulated miRNAs were found (Fig. 4E).

In an ideal transcriptomics study, the expression of at least some transcriptionally deregulated genes should correlate with the expression of some transcriptionally deregulated miRNAs. We applied the TargetScan algorithm [35] to identify matches between deregulated genes and deregulated miRNAs. A total of 1417 deregulated genes were targeted by at least one family of miRNA found deregulated in this study (Supp. Table 10). miRNAs families are composed by different miRNAs that share the same seed region [24]. The putative interactions described in Supp. Table 10 are a valuable source of hypothesis to be experimentally validated.

### Copy number variants

CNVs include amplification (gains) or deletions (losses) of chromosomal fragments and are often associated with disease [36]. We analyzed CNVs in each of our HCC samples as compared to their non-cancer counterpart. In general, amplifications were more common than deletions (Fig. 6A and Supp. Fig. 5A). In resected HCC samples, CNVs were not distributed homogenously along the genome, instead some chromosomes exhibited considerably higher frequency of CNVs, like chromosomes 1, 8, 17 and Y. Conversely, some chromosomes like 2, 3, 4, 13, 14 and 18 exhibited few CNV events. The rest of chromosomes deployed an intermediate phenotype (Fig. 6B). Hierarchical clustering of CNVs delineated three subgroups of resected tumors (Fig. 6C).

**Figure 6.**
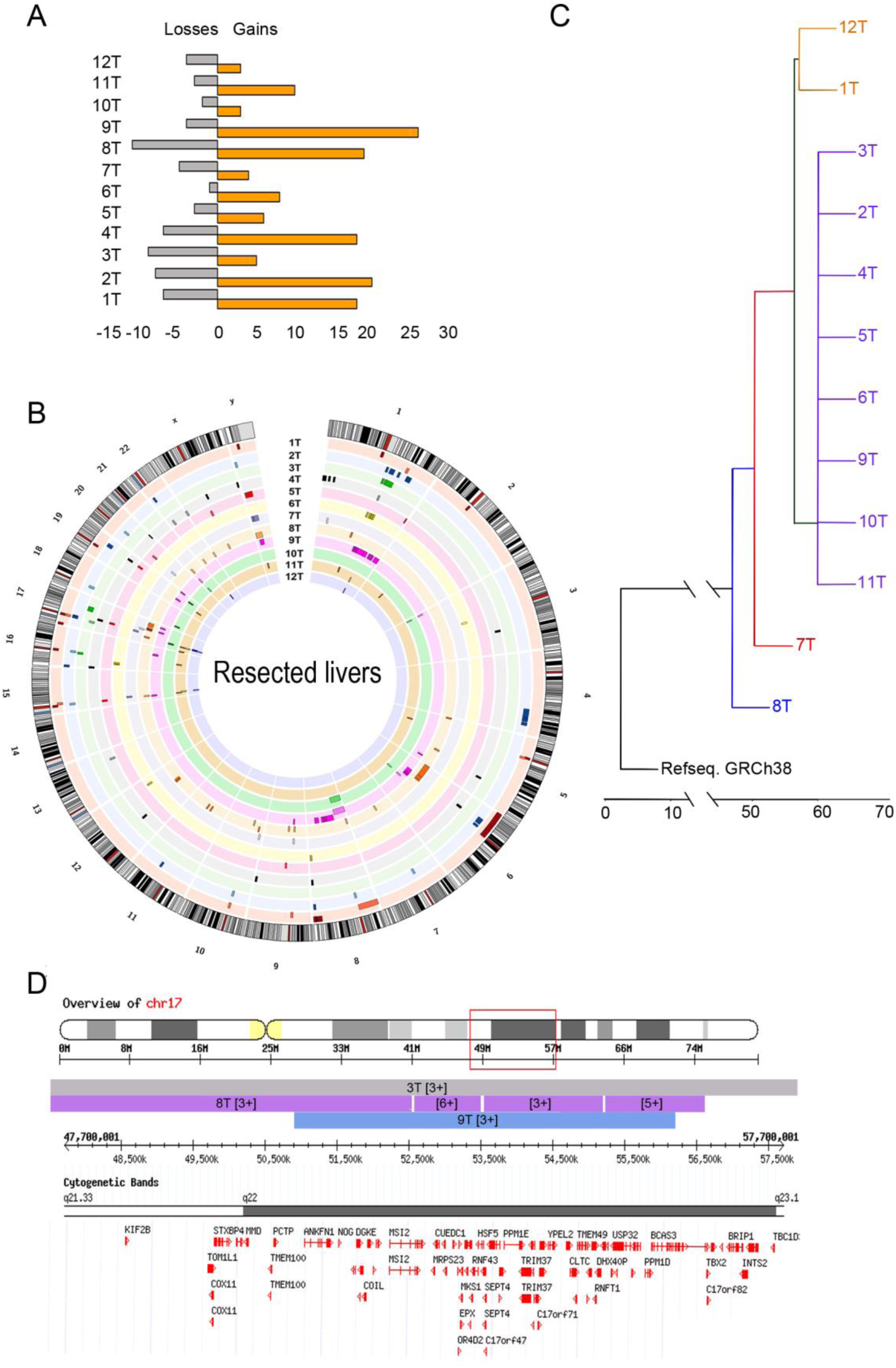
Copy number variants profile is highly variable between samples and is generally associated with amplification of oncogenes. (**A**) Number of CNV events (losses and gains) in each of the resected tumor samples. (**B**) Ideogram depicting the location and size of the CNV events in each resected tumor sample (copy number is not show here, but is shown in Supp. Table 10. (**C**) A dendrogram was derived from the CNV profiles using hierarchical clustering with complete linkage and the GRCh38.81 human genome assembly to root the dendrogram. For each tumor sample, its non-tumor counterpart was used as reference. Many of the amplified chromosomal regions contain genes potentially involved in tumorigenesis, as depicted on Fig. 7 and Supp. Fig. 6. For the sake of clarity, cytogenetic bands and genes included in the corresponding transect are shown. (**D**) An amplified region in HCC resected tumors that spans genes involved in cell proliferation and metastasis. A representative region on chromosome 17, which was amplified in tumor samples 3, 8 and 9 is shown. Genes TOM1L1, MSI2, MRPS23, RNF43 and INTS2 were found upregulated in the RNAseq libraries.

We inspected some CNVs that were present in several cancer samples, under the assumption that they were selected because conferred some advantage for cancer progression. On Fig. 6D, we depict some CNV events located on chromosome 17, between cytogenetic bands q21.33 and q23.1, which were found in tumor samples 3T, 8T and 9T. The amplification ranged from one additional copy in samples 3 and 9, to five additional copies in a sub-fragment amplified in tumor 8T. When contrasted to our RNAseq data, some of the genes contained in this transect were found upregulated, including TOM1L1 (2.35-fold), MSI2 (2.32-fold), MRPS23 (2.36-fold), RNF43 (3.26-fold) and INTS2 (2.71-fold). It is possible that other genes are also upregulated in these three samples, but were not found differentially expressed because of low level of expression in the rest of cancer samples. One and two more regions recurrently amplified on chromosome 1 and chromosome 17, respectively, are depicted in Supp. Fig. 6.

### Diversity of mutations in common cancer-related genes

We used the Accel-Amplicon 56G Oncology Panel v2 to assess mutation in genes previously reported in several cancers. Invariably, all tumors in our resected or explanted cohorts harbored mutations in the proto-oncogenes *Colony stimulating factor 1 receptor* (CSF1R) [37], the *Nucleolar phosphoprotein B23* (NPM1) [38], the GPCR-like receptor *Smoothened* (SMO) [39], the *Platelet-derived growth factor receptor alpha polypeptide* (PDGFRA) [40], the *fms-like tyrosine kinase 3* (FLT3) [41], the *fibroblast growth factor receptor3* (FGFR3) [42] as well as in the tumor suppressors *Phosphatase and tensin homologue* (PTEN)[43] and the cell cycle control protein *Tumor protein P53* (TP53) [44]. Twenty-six other genes were mutated at different frequency (Table 1).

## Discussion

The multitude of causal agents reported for HCC [7], as well as the mutagenic intrinsic ability of HBV [45], contribute to the molecular diversity found among individual tumors. Although considerable progress has been made by applying next generation sequencing technologies and bioinformatics approaches, our appreciation of the molecular heterogeneity of HCC is far from complete. Here, we contribute an additional dataset derived from small cohorts of HCC tumors from resected and explanted livers. In addition, we analyzed transcriptome data from 50 HCC tumors and paired controls from the TCGA project [27].

At the transcriptional level, a myriad of genes was found deregulated in tumors. The biological significance of deregulated genes was different in resected and explanted livers. In the former case, it indicated the emergence of drug resistance, the induction of inflammation and the increase of cell proliferation, all processes in line with tumor growth [46]. In explanted livers, a marked depletion of processes related to energy production was observed as well as moderate enrichment of functions related to hypoxia tolerance, which altogether may suggest a strong reliance of cancer cells on glycolysis for energy production [47].

Individual genes warrant further investigation. For instance, the C-type lectin CLEC4G (aka LSECtin), was found 156-fold downregulated in our study and 538-fold in the TCGA cohort, and has been reported to favor metastasis of colon carcinoma to the liver, likely by promoting adhesion of cancer cells in the liver [48]. Also, LSECtin antagonizes activated T cells in the liver, to maintain a tolerogenic environment under normal physiological conditions [49]. Thus, downregulation of LSECtin in HCC tumors may contribute to inflammation and to detachment of cells from the tumor, which will favor tumorigenesis and metastasis, respectively. The anilin actin binding protein (ANLN) was found 49-fold upregulated in our data and 15-fold upregulated in the TCGA cohort, and has been implicated in lung cancer progression [50]. Many other examples could be mentioned.

Deregulated genes also includes lncRNAs, like upregulation of LINC00152, which has been implicated in progression of gastric cancer and HCC [51–54], or downregulation of LINC01018 which, when epigenetically silenced, has been shown to promote HCC proliferation [55]. The top deregulated lncRNA in our study was LINC01093 (fold-change 30.4), which has been reported downregulated in association with HCC progression [56,57]. Thus, studying the mechanisms whereby LINC01093 antagonizes HCC might be a promising avenue for drug discovery, as it could be supplied exogenously to HCC patients or could also serve as a biomarker of disease. The rest of lncRNAs depicted in Fig. 3 have not been previously implicated in HCC and their potential contribution to disease remains to be elucidated. miRNAs are another important class of regulatory non-coding RNAs involved in liver diseases [24,25,58]. The most dramatically deregulated miRNA in our samples was miR-1269a (73-fold upregulated), which has been reported associated with colorectal carcinoma relapse and metastasis [59], relapse of esophageal squamous cell carcinoma [60], and proliferation of HCC via downregulation of FOXO1 [61]. However, we only detected an slight downregulation of FOXO1 (2.9-fold), which suggests that other targets of miR-1269a may also be involved in the HCC pathology. Another interesting example is miR-4485-3p, which was downregulated in our data, was found deregulated in a gastric cancer cell line [62], and has been suggested to counter angiogenesis in Kaposi’s sarcomas [63]. Given its therapeutic potential as an antagonist of angiogenesis, further research on this miRNA is needed.

Copy number variants were abundant and represent a mechanism to promote tumorigenesis. Especially interesting is Chr 17, because of its high susceptibility to structural variations in several of our tumors, which suggest that they have been shaped by selection pressure during cancer evolution. Some amplified regions on Chr 17 contained genes upregulated in the RNAseq data. This included MRPS23, which has been associated with metastatic phenotypes of uterine cervical cancer [64] and breast cancer [65]; MSI2, which promotes metastasis of non-small cell lung cancer through regulation of TGFβ [66] or TOM1L1, which promotes breast cancer cell invasion [67]. Finally, the tumor genomes were found highly lacerated. Out of 56 genes screened, 34 were mutated with different degree of penetrance. Typical tumor suppressors like PTEN and TP53 and few other cancer-related genes were mutated in all tumors, as often occurs in many type of cancers, but a number of other genes were mutated in some, but not all, tumors. This further highlights the molecular heterogeneity of individual HCC tumors.

A unifying theme in our data is heterogeneity across samples. Recently, it has been proposed that cancer could be better managed if they are stratified into subgroups that are molecularly and clinically similar. Clustering techniques applied to our RNAseq, miRNAseq and CNV profiling lead to substantially different subgroups of tumor samples, evidencing an intrinsic limitation of stratification of tumors based on single layers of information (i.e. genomics, transcriptomics, CNVs, etc.). A possible solution to such problem could be the incorporation of multiple omics data sets together with clinical metadata and disease outcome information, into machine learning approaches able to model the combined contribution of several layers of information and its correlation with disease outcome. This is consistent with a biological scenario where a battery of protein-coding genes, non-coding RNAs, mutant proteins, copy number variants, silent loci and other attributes act in concert to mediate transition towards the cancer phenotype. Performance of such computational approaches will increase as data sets from different laboratories across the world continue to accumulate, broadening the inventory of molecular aberrations in cancers. We expect that the information we generated in this study will elicit hypotheses that other researchers will evaluate, to ultimately increase our comprehension of liver cancer biology.

## Materials and Methods

### Samples, nucleic acids and libraries

Explanted liver samples were collected according to protocols reviewed and approved by the Human Research Ethics Board of the University of Alberta. Resected liver samples were acquired from the tumor bank at the Cross Cancer Institute, University of Alberta.

Total RNA was extracted with TRIzol Reagent (Invitrogen) as per manufacturer recommendations and the same prep was used for RNAseq and miRNA libraries construction. Genomic DNA was extracted using the DNeasy Blood & Tissue kit (QIAGEN) following manufacturer’s protocol. RNA integrity and concentration was determined using an Agilent Bioanalyzer and a chip from the RNA 6000 nano kit. DNA was quantified fluorometrically in a Qubit instrument and a dsDNA HS assay kit.

For construction of RNAseq libraries, the Illumina TrueSeq^®^ RNA Sample Preparation Kit V2 was used as per manufacturer’s instructions. Libraries from explanted livers were sequenced in a HiSeq 2500 instrument, using a paired-end 300 cycles protocol at an average depth of ~ 40 M paired end reads per sample. Samples from explanted livers were sequenced in a MiSeq instrument using a paired-end 150 cycles protocol at an average depth of ~ 2 M paired-end reads per sample. Whole genome libraries for evaluation of copy number variants were constructed using the Illumina Nextera XT library prep kit, according to manufacturer instructions. Libraries were sequenced in a MiSeq instrument using a paired-end 500 cycles kit at an average coverage of 0.15X. For detection of mutations in oncogenes the Accel-AmpliconTM 56G Oncology Panel v2 (Swift Biosciences) was used as per provided protocols. Libraries were sequenced in a MiSeq instrument using a paired-end 500 cycles kit at an average depth of 500,000 paired-end reads per sample. All samples were sequenced following a workflow that includes demultiplexing and adapter trimming.

## Bioinformatics

Low quality sequences (Q < 25) were trimmed off with the FastqMcf software and only paired-end reads with a length of at least 75% of the original length were kept for further processing.

### RNAseq libraries

Sequences were aligned to the GRCh38.81 assembly of the human genome using the TopHat2 aligner [68]. Reads that aligned to each gene sequence were counted with HTSeq [69]. Differential expression analysis was conducted with the DESeq2 R package and genes that were deregulated 2-fold or more and had a false discovery rate < 0.05 were considered truly differentially expressed.

### miRNA libraries

Libraries adapters were removed with in-house Perl scripts. Trimmed sequences were aligned to the version 21 of the miRBase mature sequences with the Bowtie aligner [70]. Aligned sequences were parsed with in-house Perl scripts and differential expression analysis was conducted as above.

### Copy Number Variants

Sequences were aligned to the GRCh38.81 assembly of the human genome using the Bowtie2 aligner [71]. Sam files were then used to infer CNVs using the copy number estimation by a mixture of PoissonS (CN-MOPS) R package [72], using paired non-tumor samples as references.

### Detection of mutations in oncogenes

Mutations were detected using the Genome Analyzer Toolkit (GATK) [73]. In brief, sequences were aligned to the g1k_v37.fasta human genome sequence provided by GATK (https://software.broadinstitute.org/gatk/) with the BWA aligner [74]. Reads were tagged, sorted and indexed with the Picard tools (http://broadinstitute.github.io/picard). Base qualities were recalibrated and indels were realigned with GATK. Variants were called with the HaplotypeCaller of GATK and finally filtered (FS > 30; QD < 2.0) with the VariantFiltration program of GATK. All plots were created using the R programing language or Circos [75]. Parsing and pre-processing of data was carried out with in-house Perl or Python scripts.

---

**Supplementary Figure 1.**
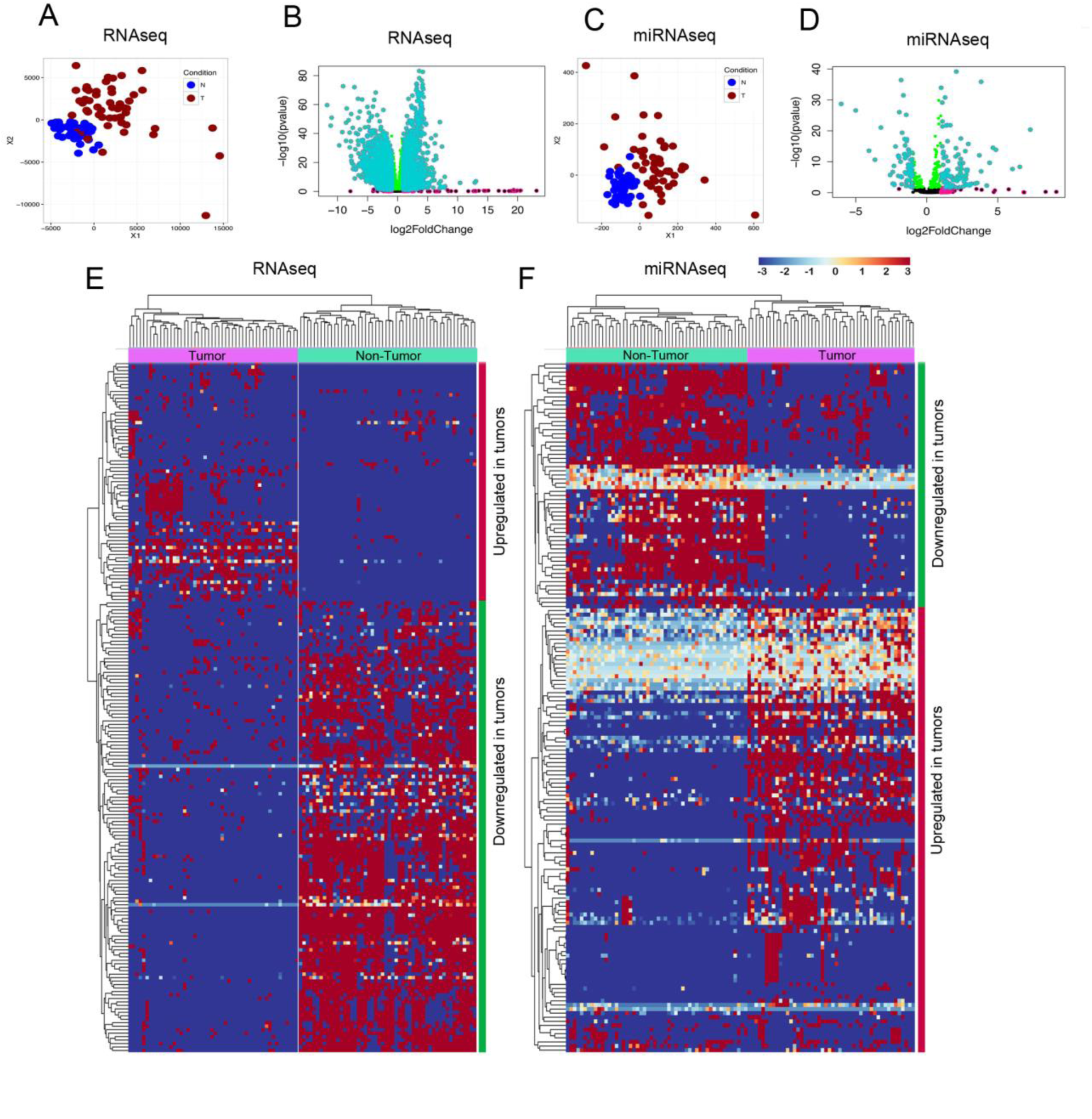
RNAseq and miRNA profiles for the TCGA cohort. Data was downloaded from the TCGA data portal (https://tcga-data.nci.nih.gov/docs/publications/tcga/?) and then processed using the same pipeline we used to analyze our own data. Only 50 tumros for which paired non-tumor controls were available were included in this analysis. (**A**) Principal component analysis for RNAseq data. (**B**) Differentially expressed genes (**C**) Principal component analysis for miRNAs data. (**D**) Differentially expressed miRNAs. (**E**) Hierarchical clustering of top differentially expressed genes. (**F**) Hierarchical clustering of top differentially expressed miRNAs.

**Supplementary Figure 2.**
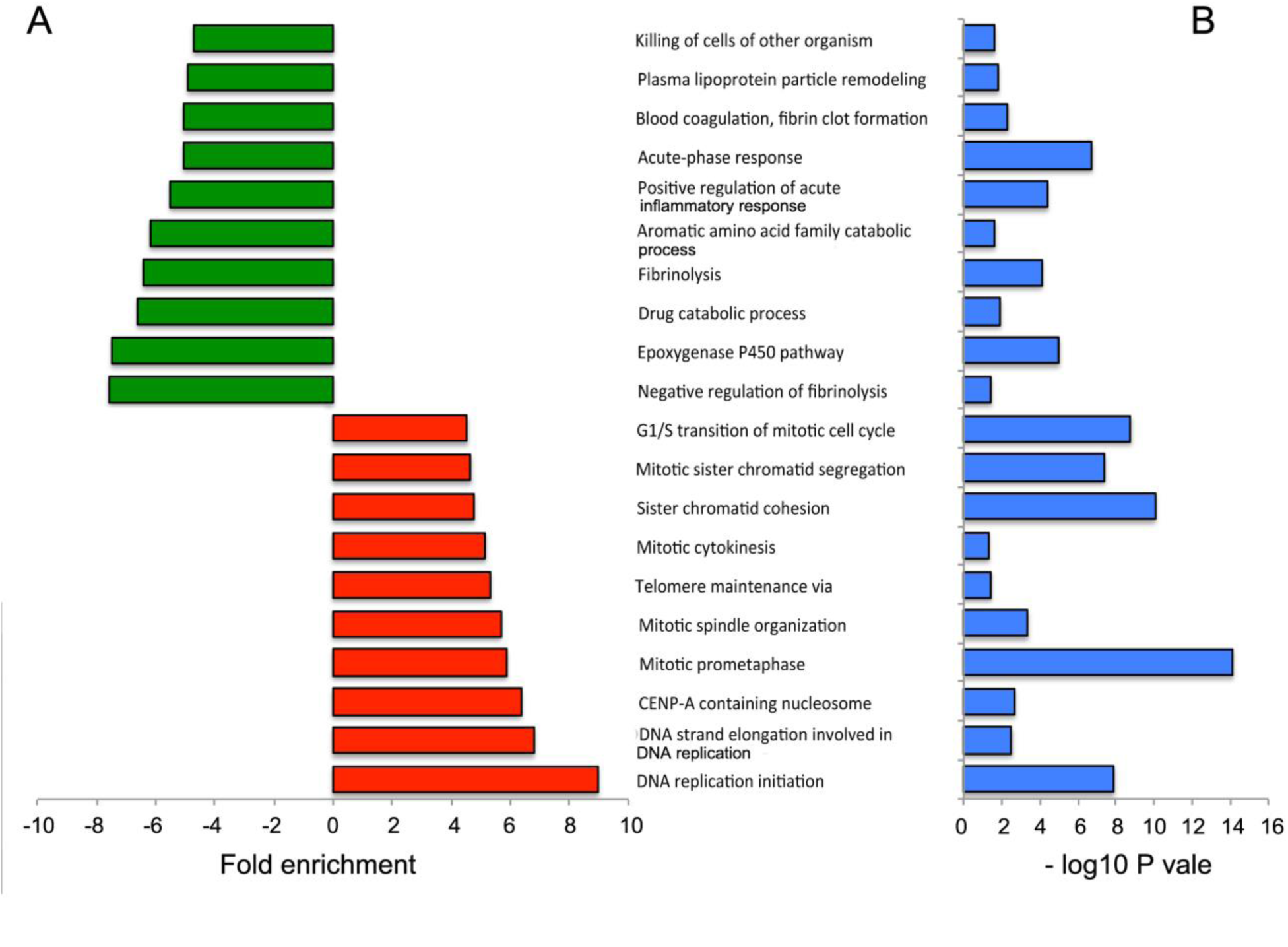
Gene ontology analysis of the TCGA cohort. As in the case of our resected livers cohort, the predominant deregulated gene ontology terms in the TCGA cohort were related with activation of the immune systems and emergence of drug resistance. Cell proliferation and DNA synthesis were the most enriched terms.

**Supplementary Figure 3.**
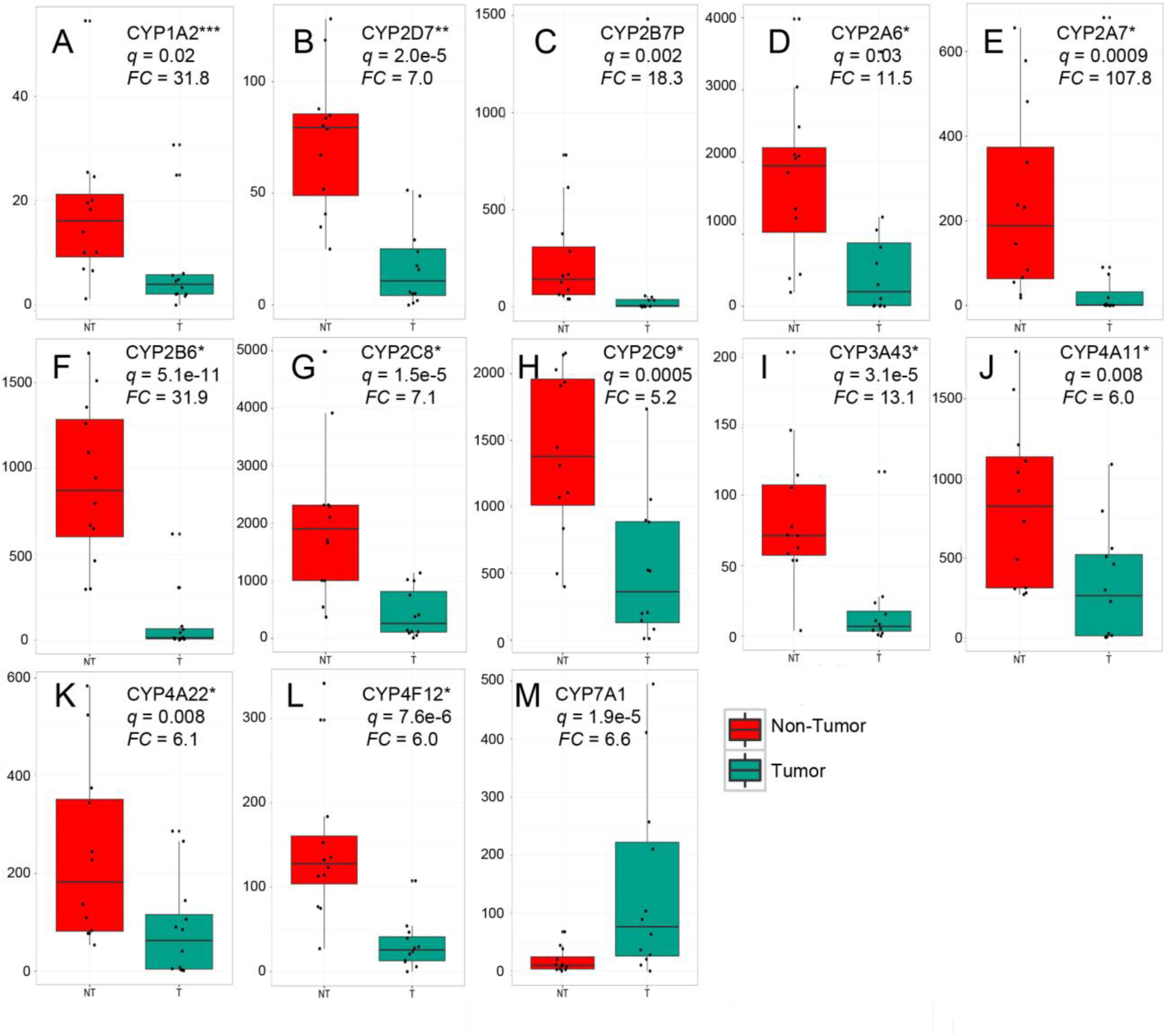
Many monooxigenases in the cytochrome P450 superfamily are deregulated in resected HCC tumors. Upregulated genes included the monooxygenases (**A, B, C**), and oxidase involved in oxidation of nicotine of cotidine (**D, F**), oxidases involved in the metabolism of long-chain polyunsaturated fatty acids (**G**), oxidases involved in metabolism of arachidonic acid and about 100 drugs (**H, J, K**), enzymes involved in the synthesis of cholesterol, steroids and other lipids (**L**). The only oxygenase found upregulated was the cholesterol 7 alpha hydroxylase (CYP7A1) (**M**). *: found deregulated in our resected cohort and in the TCGA cohort. **: Found deregulated in our resected and explanted cohorts. ***: found deregulated in the three cohorts.

**Supplementary Figure 4.**
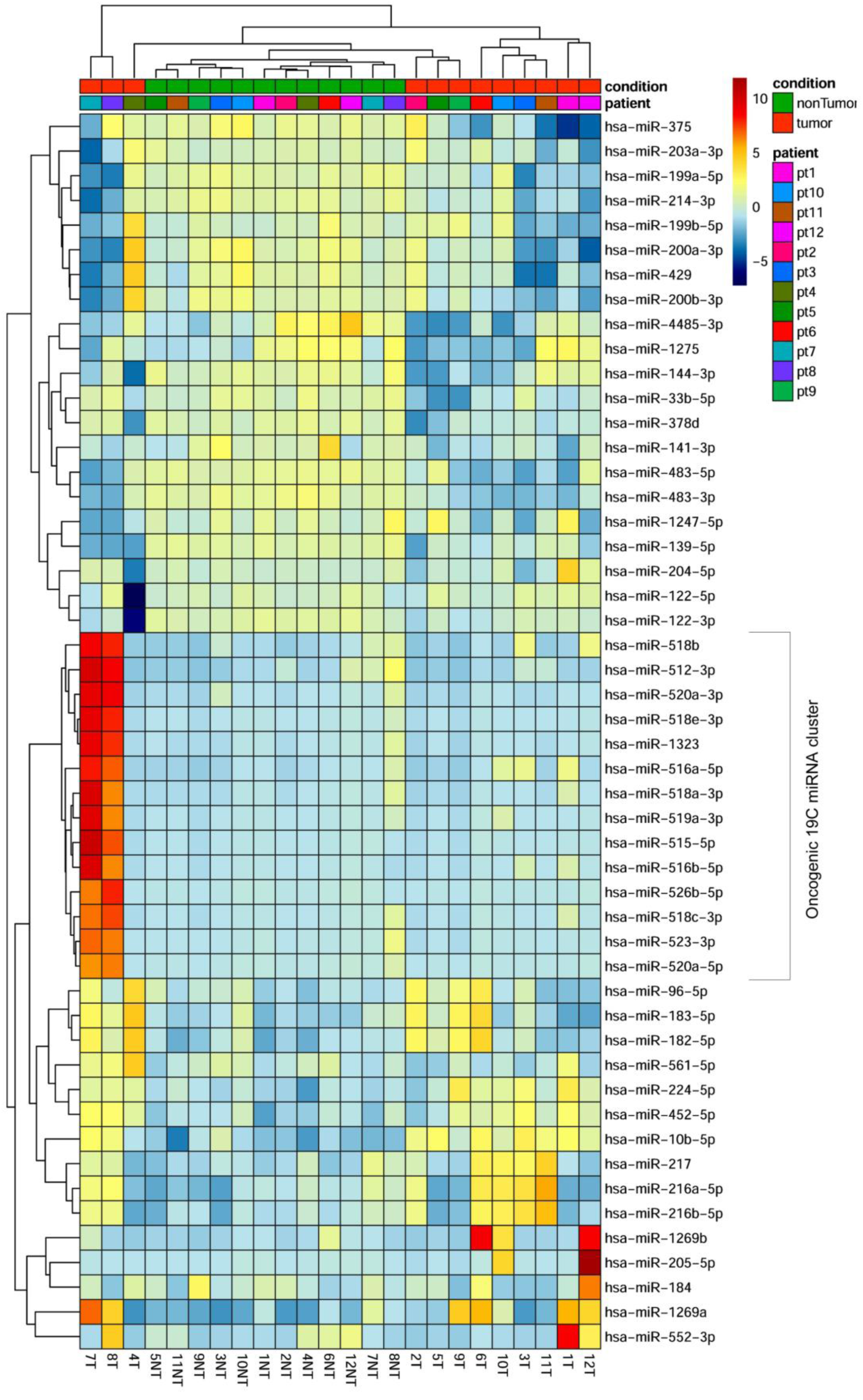
Tumor samples 7 and 8 exhibit a dramatic upregulation of miRNAs in the 19C cluster. Hierarchical clustering of 50 miRNAs exhibiting the largest variance between groups(cancer vs non-cancer), irrespectively of whether they were differentially expressed or not, illustrates a dramatic upregulation of many miRNAs in the 19C cluster in tumors 7 and 8. This may explain the separation of these two samples in the principal component analysis plot presented in Fig. 5A.

**Supplementary Figure 5.**
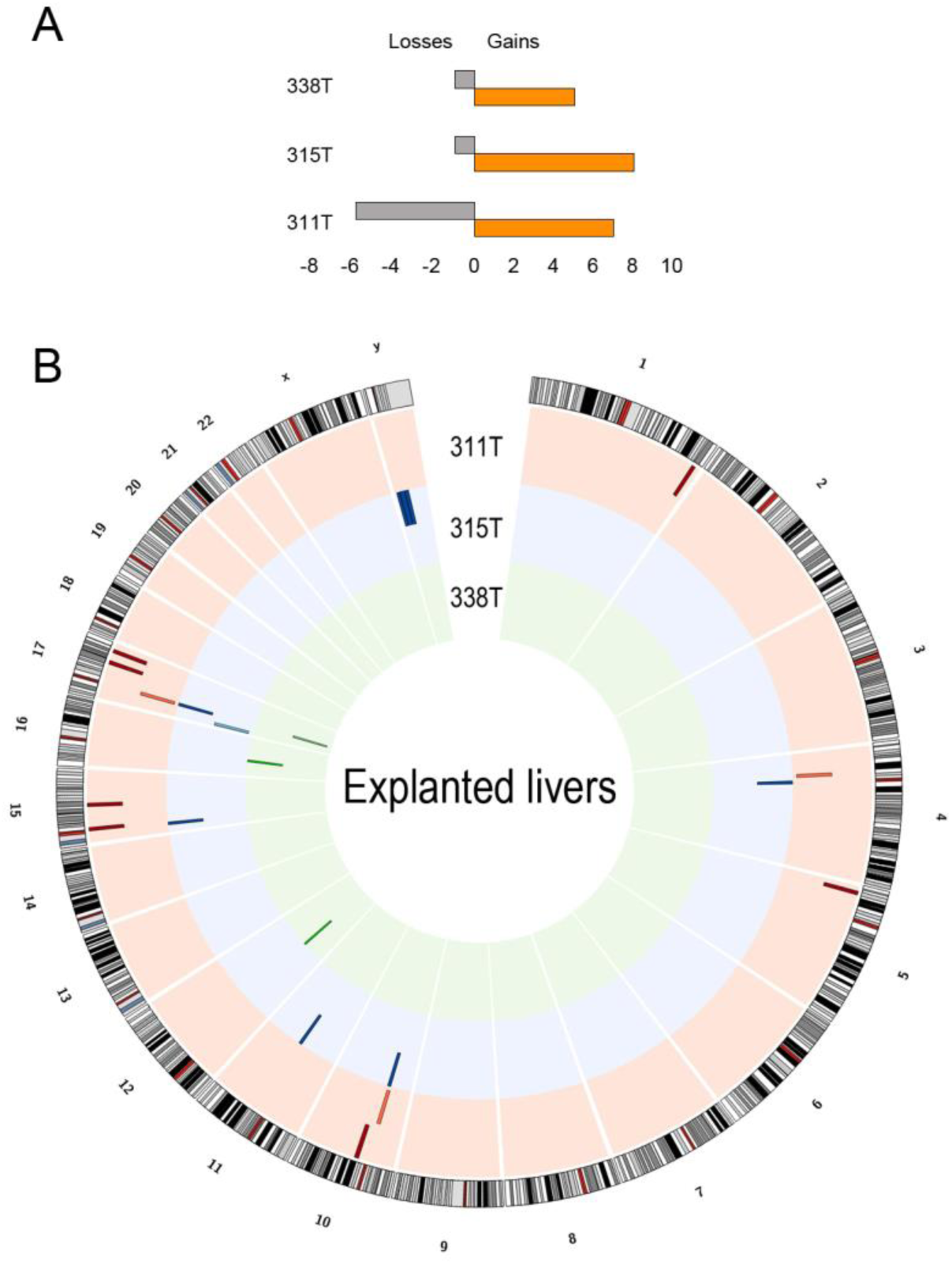
Copy number variants profile of tumors in explanted livers. (**A**) As inthe case of resected livers (Fig. 6), number of CNV events demonstrate that gains were more common than losses in explanted livers. (**B**) CNV events were not distributed evenly along all chromosomes of the genome. Chromosome 17 seems to constitute a hotspot for CNV events generation. In general, explanted livers presented less CNV events than resected livers. The reasons for such difference is not known.

**Supplementary Figure 6.**
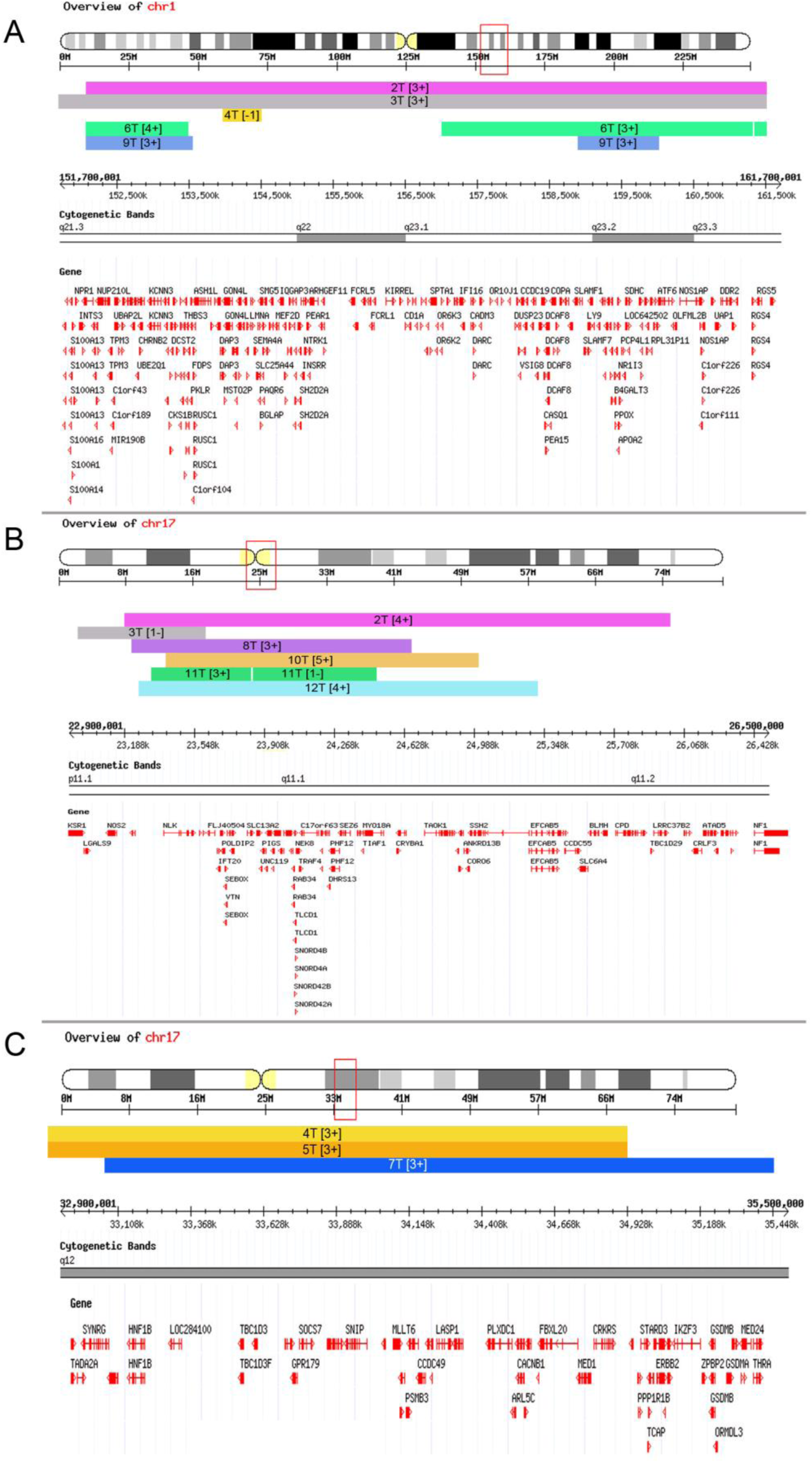
Recurrent CNV hotspots found in chromosomes 1 or chromosome 17. For the sake of clarity, cytogenetic bands and genes included in the corresponding transect are shown

**Supplementary Table 1.** Description of resected and explanted HCC samples analyzed in this study.

**Supplementary Table 2.** Differential expression analysis results from RNAseq data generated from 12 HCC resected livers. Downregulated genes are in blue font.

**Supplementary Table 3.** Differential expression analysis results from RNAseq data generated from 3 HCC explanted livers. Downregulated genes are in blue font.

**Supplementary Table 4.** Differential expression analysis results from microRNAseq data generated from 12 HCC resected livers. Downregulated miRNAs are in blue font.

**Supplementary Table 5.** Differential expression analysis results from microRNAseq data generated from 3 HCC explanted livers. Downregulated miRNAs are in blue font.

**Supplementary Table 6.** Differential expression analysis results from RNAseq data retrieved from TCGA corresponding to 50 HCC cancer-nonCancer pairs. Downregulated genes are in blue font.

**Supplementary Table 7.** Differential expression analysis results from microRNAseq data retrieved from TCGA corresponding to 50 HCC cancer-nonCancer pairs. Downregulated miRNAs are in blue font.

**Supplementary Table 8.** Number of genes that were found deregulated in one, two or three cohorts of samples.

**Supplementary Table 9.** HGCN symbol of genes that were found deregulated in one, two or three cohorts of samples.

**Supplementary Table 10.**Putative targets identified by TargetScan for all miRNAs found deregulated in our resected livers cohort. Targets are restricted to genes found deregulated in the RNAseq libraries from resected livers.

**Supplementary Table 11.** Copy number variants in each tumor sample of our resected and explanted livers cohorts, as determined by cn-mops.

**Supplementary Table 12.** Allelic frequency of mutations in genes related to cancer according to the Accel-Amplicon 56G oncology panel v2. The number of mutations in each gene is not necessarily an indicative of the actual mutation susceptibility because the number of amplicons per gene in the assay used is variable.

